# A scalable cell-free manufacturing platform for two-step bioproduction of immunogenic conjugate vaccines

**DOI:** 10.1101/2025.08.05.668792

**Authors:** Derek A. Wong, Rochelle Aw, Sophia W. Hulbert, Yufan Qin, Zachary M. Shaver, Kathryn A. Myers, Ashty S. Karim, Matthew P. DeLisa, Michael C. Jewett

## Abstract

Rapid and decentralized vaccine production is essential to ensure global preparedness against emerging and re-emerging infectious diseases. Cell-free gene expression systems, which can be freeze-dried for long-term storage and re-activated for point-of-use synthesis, offer a promising solution to address this need. However, scalable cell-free production of conjugate vaccines—highly effective tools against bacterial infections—has been hindered by low yields and inefficient glycosylation. Here, we present an optimized, modular cell-free platform for the synthesis and purification of conjugate vaccines. By decoupling cell-free protein expression from *in vitro* glycosylation in a two-step approach, we achieve >85% glycosylation efficiency and up to ∼450 mg/L of glycoprotein. We apply this platform to manufacturing protein-polysaccharide conjugates composed of vaccine carrier proteins covalently modified with polysaccharide antigens from enterotoxigenic *Escherichia coli* O78 and *Streptococcus pneumoniae* serotype 4. Our workflow produced conjugate vaccine candidates in under 5 days with >87% product purity and low endotoxin levels suitable for preclinical evaluation. Immunization of mice with the pneumococcal conjugate vaccine induced a strong IgG response against the *S. pneumoniae* serotype 4 capsular polysaccharide, confirming the immunogenicity of the conjugate. We anticipate that this cell-free platform will advance efforts in decentralized manufacturing and rapid response to bacterial disease threats.

## Introduction

Bacterial infections remain a significant global health challenge, especially as antibiotic resistance increases. Vaccines are among the most effective tools for preventing bacterial diseases, as they reduce both the incidence of infection and the need for antibiotic treatment, which helps limit the spread of resistant strains (1). Unfortunately, most protein-based countermeasures against bacterial pathogens are currently produced in centralized facilities with months-long lead times, expensive production steps, and the requirement of cold chain storage conditions for distribution (2).

Cell-free gene expression (CFE) systems have emerged as a technology to address these limitations. These systems harness the cellular machinery in crude lysates to enable biological processes, including protein synthesis, in open and tunable reaction environments (3–8). Owing to improvements in cell-free technology over the last two decades, protein yields from cell-free reactions exceed gram protein per liter reaction volume, batch reactions last for hours, and reaction scales have reached the 100-liter milestone (9). Furthermore, these reactions may be freeze-dried for long-term, non-refrigerated storage and transported to the point-of-care, enabling the decentralized manufacturing of therapeutics and vaccines (8, 10–14). These advantages place cell-free systems at the forefront of protein biosynthesis technology with implications for medicines (15), diagnostics (16–25), metabolic engineering (26–28), and educational kits (29–34).

Recently, CFE systems have been adapted to synthesize a wide variety of glycoproteins, including several with therapeutic relevance (10, 13, 35–44). In one such system, protein glycan coupling technology, which utilizes oligosaccharyltransferases (OSTs) to site-specifically transfer membrane bound glycans onto a protein, has been adapted to produce conjugate vaccines (45). These vaccines, which consist of a pathogen-specific O-polysaccharide (O-PS) or capsular polysaccharide (CPS) antigen attached to an immunogenic carrier protein, promote a robust anti-pathogen immune response, and are effective tools to prevent bacterial infections (1, 46) Thus far, cell-free systems have been used to synthesize conjugate vaccines that target two different bacterial pathogens, enterotoxigenic *Escherichia coli* (ETEC) serotype O78 and *Francisella tularensis* subsp. *tularensis* Schu S4 (10, 13, 36). However, most cell-free conjugate vaccine development has been conducted using small reaction volumes (15 μL scale) that are characterized by relatively low protein yields (<200 mg/L). Additionally, these systems have only been interfaced with basic gravity flow purification methods which have resulted in low purity (<50% glycosylated protein). Robust procedures to scale-up vaccine synthesis and purify glycoconjugate products produced in CFE systems have yet to be optimized, thereby limiting the utility of CFE for scalable, rapid production of glycoconjugate proteins that can be evaluated preclinically, including animal studies.

In this work, we set out to address this gap and improve the yield and purity of cell-free produced conjugate vaccines. First, we optimized reaction conditions to synthesize protein-polysaccharide conjugates composed of protein D (PD) from *Haemophilus influenzae*, a carrier protein used in commercial vaccines such as Synflorix (47) conjugated with either the O-PS from ETEC O78, a cause of diarrheal disease worldwide (48), or CPS from *Streptococcus pneumoniae* serotype 4 (CPS4), which can cause pneumonia (49). The key design choice was to separate carrier protein production from *in vitro* glycosylation by the *Campylobacter jejuni* OST, PglB (*Cj*PglB). Through these optimizations, we achieved maximum glycosylation efficiencies of >85% and vaccine yields of ∼450 mg/L in unpurified reactions. Next, we scaled our reactions three orders of magnitude and optimized downstream purification, enrichment, and endotoxin removal steps (**Figure 1**). These steps were used to produce 16 purified doses of PD conjugated to ETEC O78 O-PS (PD-O78) and 40 purified doses of PD conjugated to CPS4 (PD-CPS4) with >87% purity in 5 days from 7.5 and 30 mL *in vitro* glycosylation reactions, respectively. Finally, we demonstrated that our PD-CPS4 vaccine produced a robust CPS4-specific IgG antibody response in mice. This work paves the way for the distributed manufacturing of high-yield conjugate vaccines using CFE systems that elicit pathogen-specific immune responses.

**Figure 1.**
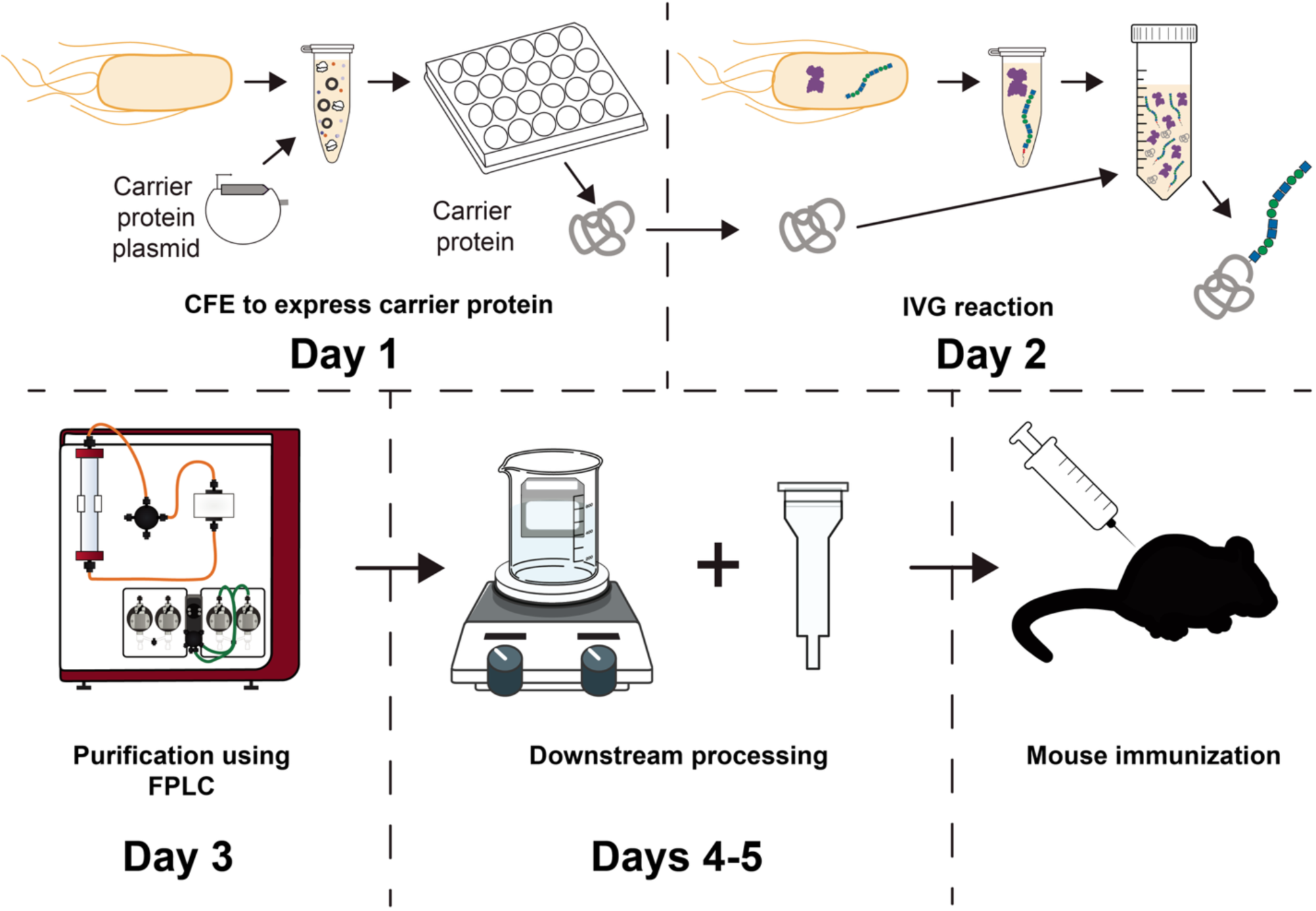
Scalable two-step cell-free production workflow for conjugate vaccines. Schematic representation of the *in vitro* glycosylation-based workflow used for conjugate vaccine production, including associated time frames for each step. FPLC: Fast Protein Liquid Chromatography.

## Results

### Assessing one-step versus two-step cell-free production of conjugate vaccines

We set out to establish a robust, high-yielding, and scalable *in vitro* method for producing conjugate vaccines. Previously, we established a one-pot cell-free glycoprotein synthesis (CFGpS) method (10, 13, 36, 50) (**Figure 2A**). For CFGpS, both the biosynthesis pathway for a bacterial glycan of interest and an OST are overexpressed in *E. coli*. These cells are then used to produce an extract enriched with glycosylation machinery, which can subsequently be used to express a vaccine carrier protein in a CFE reaction. Unfortunately, the cells exhibit slow growth rates while over-expressing glycosylation machinery, which includes large glycan biosynthesis pathways and membrane-bound OSTs (**Supplementary Figure 1**). We wondered if this could lead to reduced ability to produce proteins, since cell-free protein synthesis activity is lower when growth rates are lower (51). We hypothesized that separating, or decoupling, protein production from *in vitro* glycosylation (IVG) (**Figure 2B**) could improve system performance and glycoprotein production.

**Figure 2.**
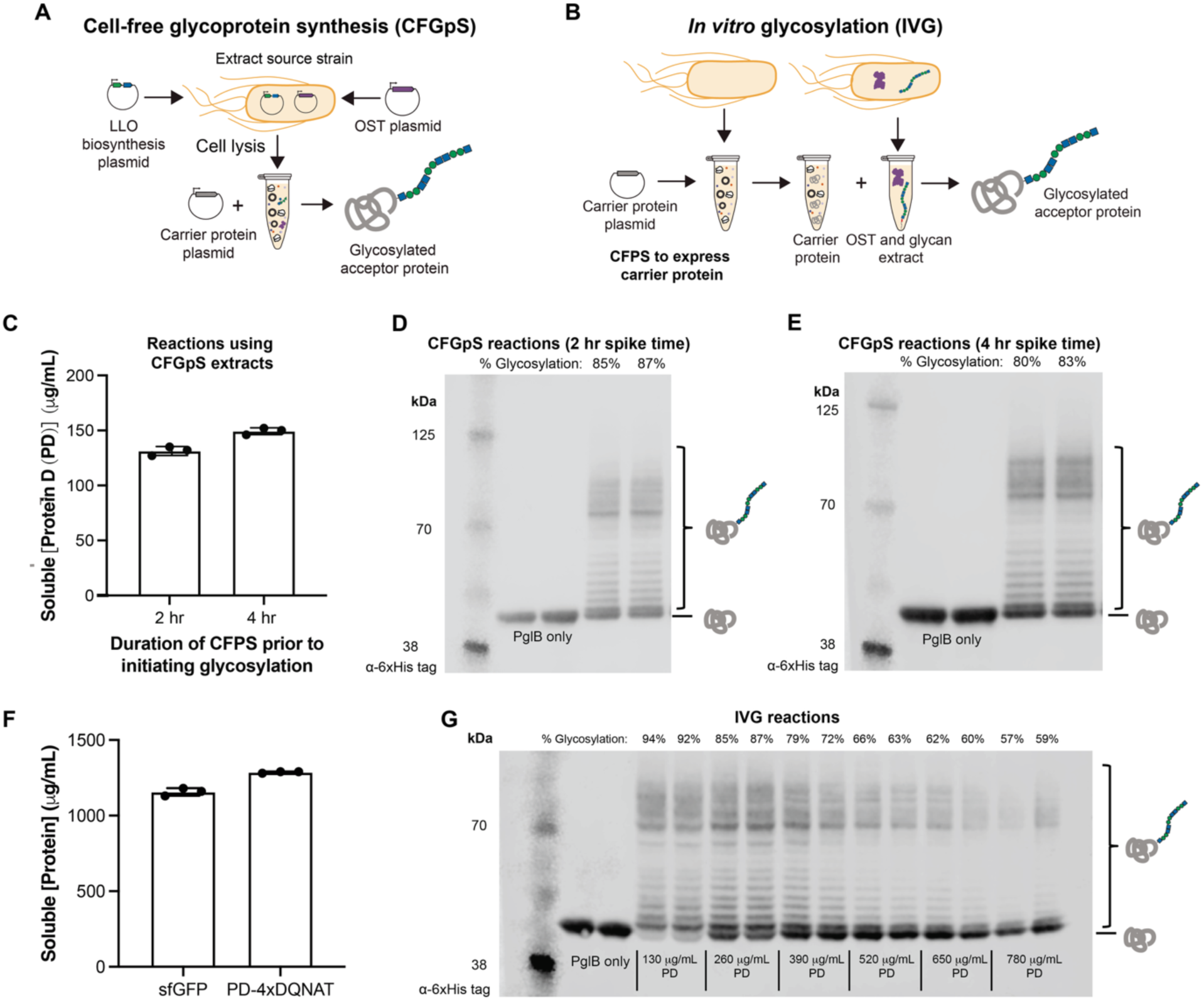
Decoupling protein synthesis and glycosylation reactions increases yields of glycoconjugate vaccines in CFE reactions. (A) Schematic of cell-free glycoprotein synthesis (CFGpS) reactions. Both lipid-linked oligosaccharide (LLO) biosynthesis pathway and OST are overexpressed during cell growth to produce an extract enriched with glycosylation machinery. The resulting extract is then used to produce a carrier protein and initiate a subsequent glycosylation reaction. (B) *In vitro* glycosylation (IVG) reactions decouple CFE and glycosylation reactions by first expressing a carrier protein in a CFE reaction and subsequently mixing with extracts enriched with glycosylation machinery. (C) Yields of PD, quantified via ^14^C-leucine, expressed in CFGpS reactions utilizing extracts enriched with *Cj*PglB and ETEC O78 O-PS. Reactions were spiked with glycosylation cofactors after either 2 or 4 h of protein synthesis. Data shown are the average of three (*n*=3) biological replicates. CFGpS reactions spiked with glycosylation cofactors after (D) 2 h or (E) 4 h of protein synthesis achieved high levels of glycosylation. An anti-6xHis Western blot was used to detect the 6xHis-tag included on the PD construct and densitometry was used to estimate percent glycosylation efficiency. Data shown are two (*n*=2) biological replicates of each condition. (F) Yields of sfGFP and PD, quantified via ^14^C-leucine, expressed in CFE reaction utilizing BL21 (DE3) Star extracts. Data shown are the average of three (*n*=3) biological replicates. (G) IVG reactions were assembled with increasing amounts of PD expressed in CFE reactions and analyzed for glycosylation efficiency using an anti-6xHis Western blot. Glycosylation efficiency was estimated using densitometry, with each condition run in sets of two (*n*=2) biological replicates.

To assess our hypothesis, we first characterized reactions using extracts enriched with the *Cj*PglB OST and O-PS from ETEC serotype O78, which we have previously used to produce immunogenic conjugates of PD-O78 (10, 36). Following protein synthesis initiation, the CFE reaction was spiked with manganese and n-dodecyl-ß-D-malopyranoside (DDM) to enable site-specific glycosylation of the expressed carrier protein by *Cj*PglB, all in a single pot. We used ^14^C-leucine incorporation to quantify the amount of protein produced during the CFE reaction prior to initiating glycosylation (**Figure 2C**). Following 2 and 4 h of protein expression, we were able to produce up to 149 μg/mL of PD, which was ∼85% glycosylated, calculated as the glycosylated bands divided by the sum of the glycosylated and aglycosylated bands (**Figure 2D & 2E**).

To test whether we could improve yields of our glycoconjugates by decoupling protein synthesis from overexpression of glycosylation machinery, we implemented a two-step IVG approach (**Figure 2B**). This system uses a high yielding CFE extract produced from BL21 (DE3) Star to produce the carrier protein. Following protein expression, the CFE reaction is then mixed with an enriched extract containing the OST and glycan of interest to enable glycosylation of the carrier protein. Using BL21 (DE3) Star derived CFE extract, we were able to produce 1287 ± 5.7 μg/mL of PD (**Figure 2F**), approximately 9-fold higher than was obtained using CFGpS reactions. Furthermore, when we use the CFE expressed PD in a subsequent IVG reaction with *Cj*PglB, we were able to obtain more glycosylated product (**Figure 2G**) when compared to the one-pot CFGpS. Specifically, using the lowest amount of PD in the IVG reaction, we obtained up to ∼93% glycosylation efficiency. Although the percentage of glycosylated protein decreased as the amount of starting protein increased, the amount of total glycoprotein produced in the reaction increased with each increment to a maximum of ∼450 μg/mL, approximately 3.7-fold higher than achieved in CFGpS. We also investigated an alternative IVG reaction set-up in which we mixed the PD-producing CFE reaction with separately enriched extracts overexpressing either *Cj*PglB or ETEC O78 O-PS (**Supplementary Figure 2A-B**). With this alternative set-up, we observed lower glycosylation efficiency compared to using a single co-enriched glycosylation competent extract (**Supplementary Figure 2C**).

### Scale-up of conjugate vaccine synthesis

To evaluate the scalability of our system, we aimed to make at least 100 µg of PD-O78 glycoconjugates (10 doses, each 10 µg) within a week with purity and glycosylation efficiency each at >80%. This process was informed by our previous purification of T7 RNA polymerase using strep-tag purification and endotoxin removal (15). Four CFE reactions of 250 µL were incubated in 24-well plates for 16 h at 30 °C. Following protein expression, a 7.5 mL IVG reaction was then set up using the conditions that yielded the highest glycosylation efficiency in our previous optimization experiment (10% v/v CFE reaction expressing PD, 80% v/v extract co-enriched for both ETEC O78 O-PS and *Cj*PglB, 10% v/v spiking solution containing manganese, Ficoll-400, HEPES buffer, and DDM). We then purified our PD-O78 glycoconjugates using a 1 mL Strep-Tactin XT column with an ÄKTA Avant and analyzed the elution fractions using SDS-PAGE to select for fractions that contained our PD-O78 glycoconjugate (**Supplementary Figure 3**). To reduce the presence of endotoxins, we used the *E. coli* strain CLM24 Δ*lpx*M to create our coenriched lysates containing the OST and glycan. This strain has a detoxified lipid A molecule through strain engineering that we have previously described (52, 53). Additionally, in the plasmid that encodes for the OST, we also overexpress the *F. tularensis* phosphatase *LpxE* gene that has been shown to lead to reduced toxicity but retained adjuvant activity (13, 52). Previously, glycoconjugate vaccines produced using this strain reduced endotoxin levels to below 1000 EU/mL. However, we incorporated an endotoxin removal step to establish a more stringent workflow. Therefore, after pooling all fractions containing our PD-O78 glycoconjugate, we used a commercially available endotoxin removal resin.

Following endotoxin removal, we used anti-6xHis Western blot densitometry to determine the percent glycosylation efficiency of our purified PD product, observing greater than 86% glycosylation of both elution fractions (**Figure 3A**). Using an anti-ETEC O78 antibody, we also confirmed the identity of glycan on our glycoconjugates (**Figure 3B**). We ran our samples in an SDS-PAGE gel and stained with Coomassie blue to assess the effectiveness of our Streptactin-based purification; using densitometry, we estimate a purity of ∼96% (**Figure 3C**). Finally, using a commercially available endotoxin quantification kit, we measured the levels of endotoxin both pre-and post-purification of our sample. We achieved average endotoxin levels of 2.91 EU per dose of 10 µg of the vaccine (**Supplementary Figure 4**), which was a ten-fold reduction prior to endotoxin removal, and is below the standard in therapeutic conjugate vaccines of 12 EU per 10 µg dose (54, 55). In total, we produced 166 µg of protein-polysaccharide conjugates, which was equivalent to 16 doses of a 10 µg dose, considered a standard dose for conjugate vaccines (56, 57) (**Table 1**). Due to the speed and capability of our conjugate vaccine production platform, we were able to achieve all vaccine synthesis, purification, and analysis steps in less than one week.

**Figure 3:**
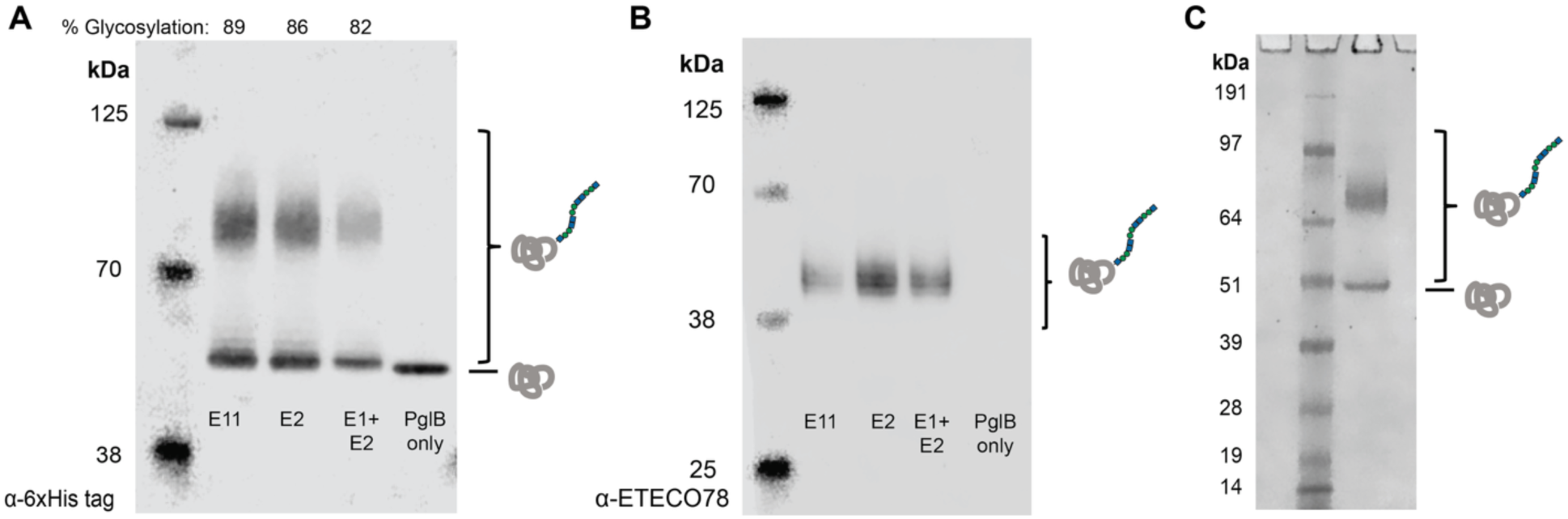
Scaled production of PD-O78 glycoconjugates in *IVG* reactions. (A) Anti-6xHis tag Western blot confirms glycosylation of PD-O78 conjugates purified from large-scale IVG reactions. (B) Anti-ETEC O78 Western blot confirms identity of glycan on PD-O78 conjugates. (C) Assessment of sample purity using a Coomassie blue stained SDS-PAGE gel of elution 1 and elution 2 mixed sample.

**Table 1:** Summary of final conjugate vaccine yields.

|  | **ETEC-O78** | **CPS4** |
| --- | --- | --- |
| **Purified conjugate yield** | 18.5 µg PD-O78 /<br>mL of IVG reaction | 8.2 µg PD-CPS4 /<br>mL of IVG reaction |
| **Purity (%)** | 96% | 87% |
| **Glycosylation (%)** | 87% | 70% |

### Production of S. pneumoniae CPS4 glycoconjugate vaccine

Once we had an established workflow that could produce quality conjugate vaccine at a high purity, we next sought to synthesize and scale up production of a therapeutic candidate targeting *S. pneumoniae* serotype 4 (CPS4), a leading cause of invasive pneumococcal disease in disadvantaged populations (49, 58, 59). PD-CPS4 glycoconjugates have previously been made in small-scale CFE reactions (60) but not yet at a scale sufficient for mouse immunization studies. To begin, we adapted our IVG reaction set-up by using extracts produced from Hobby cells overexpressing CPS4 and a mutant *Cj*PglB harboring a Q287K substitution. The Hobby strain has previously been genetically modified to produce greater amounts of CPS4 *in vivo* (61), while *Cj*PglB^Q287K^ was previously engineered by our groups to increase the transfer efficiency of CPS4 relative to wild-type *Cj*PglB (60). Using small-scale reactions, we again tested various ratios of carrier protein and glycosylation enriched extract added to the IVG reaction (**Supplementary Figure 5**). While slightly lower efficiencies were observed compared to ETEC O78 O-PS reactions, we were able to achieve ∼80% glycosylation efficiency with CPS4 at lower concentrations of PD in the IVG reaction and a maximum of ∼325 μg/mL PD-CPS4 at higher concentrations of PD.

To produce enough sample for downstream immunogenicity evaluation using a mouse study, we scaled up to a 30 mL IVG reaction using the same ratios of CFE expressed PD and glycosylation enriched extract as was used for scaling PD-O78 production. Following purification, we dialyzed our PD-CPS4 glycoconjugates into sterile PBS (pH 7.4). From a 30 mL IVG reaction we purified 400 µg of PD, which we estimated to be ∼70% glycosylated by densitometry (**Figure 4A**, **Table 1**). The identity of the glycan was confirmed using an anti-CPS4 antibody (**Figure 4B**). Despite reduced glycosylation efficiency, the purity of our product determined by SDS-PAGE was still approximately 87% (**Figure 4C)**. While we note lower glycosylation efficiency compared to our PD-O78 glycoconjugates, our glycosylation was still on par with immunogenic glycoconjugates previously produced in CFE (13, 36). As a result, our workflow was able to produce 40 10-μg purified doses in less than one week (**Figure 1**).

**Figure 4:**
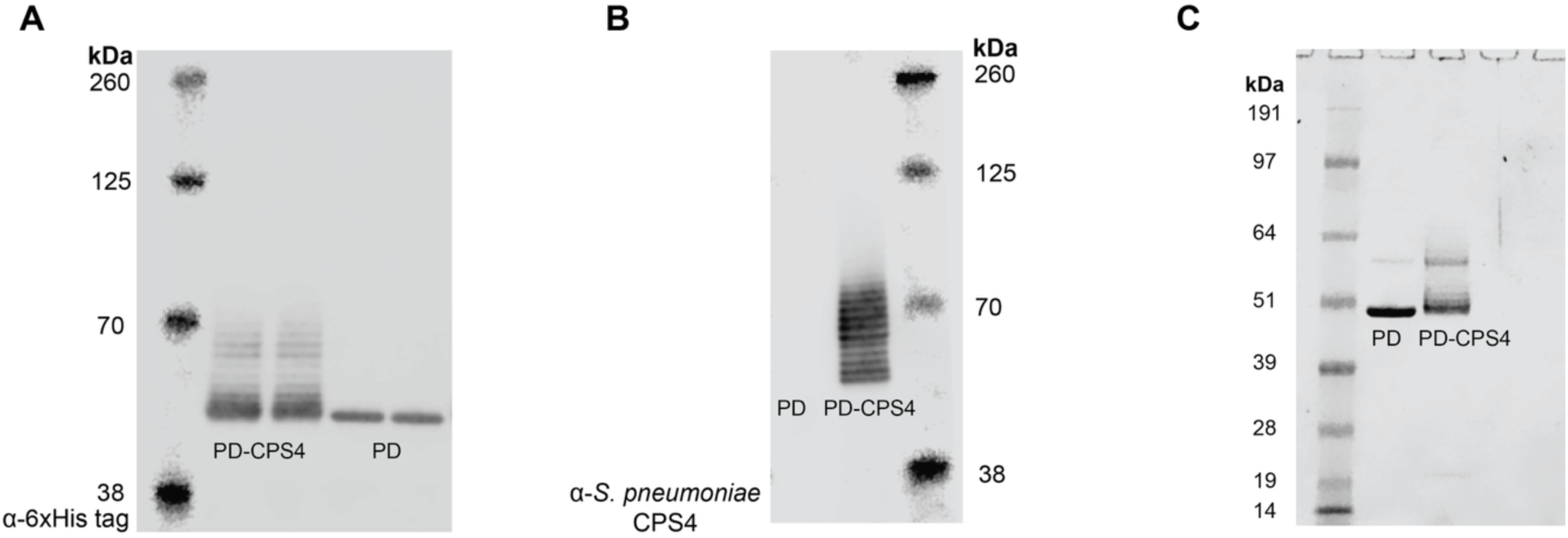
Scaled production of PD-CPS4 conjugates in IVG reactions. (A) Anti-6xHis tag Western blot confirms glycosylation of PD-CPS4 conjugates purified from one batch of large-scale IVG reactions. (B) Anti-*S. pneumoniae* CPS4 Western blot confirms identity of glycan on PD-CPS4 conjugates. (C) Assessment of sample purity of both aglycosylated PD and PD-CPS4 using an SDS-PAGE gel stained with Coomassie blue.

### In vitro synthesized PD-CPS4 conjugate elicits pathogen-specific antibodies in mice

We next evaluated the ability of our *in vitro* synthesized PD-CPS4 conjugate to elicit anti-CPS4 antibodies in mice. To this end, groups of eight mice were injected with 10 µg of the PD-CPS4 conjugate, unmodified PD carrier protein, or phosphate-buffered saline (PBS). After each dose, production of CPS4-specific IgG antibodies was monitored by ELISA. As expected, mice immunized with PD-CPS4 generated high titers of CPS4-specific serum IgG antibodies in endpoint (day 77) serum of individual mice, which were >2 logs higher than those in control mice receiving either the unmodified PD carrier or PBS (**Figure 5A**).

**Figure 5.**
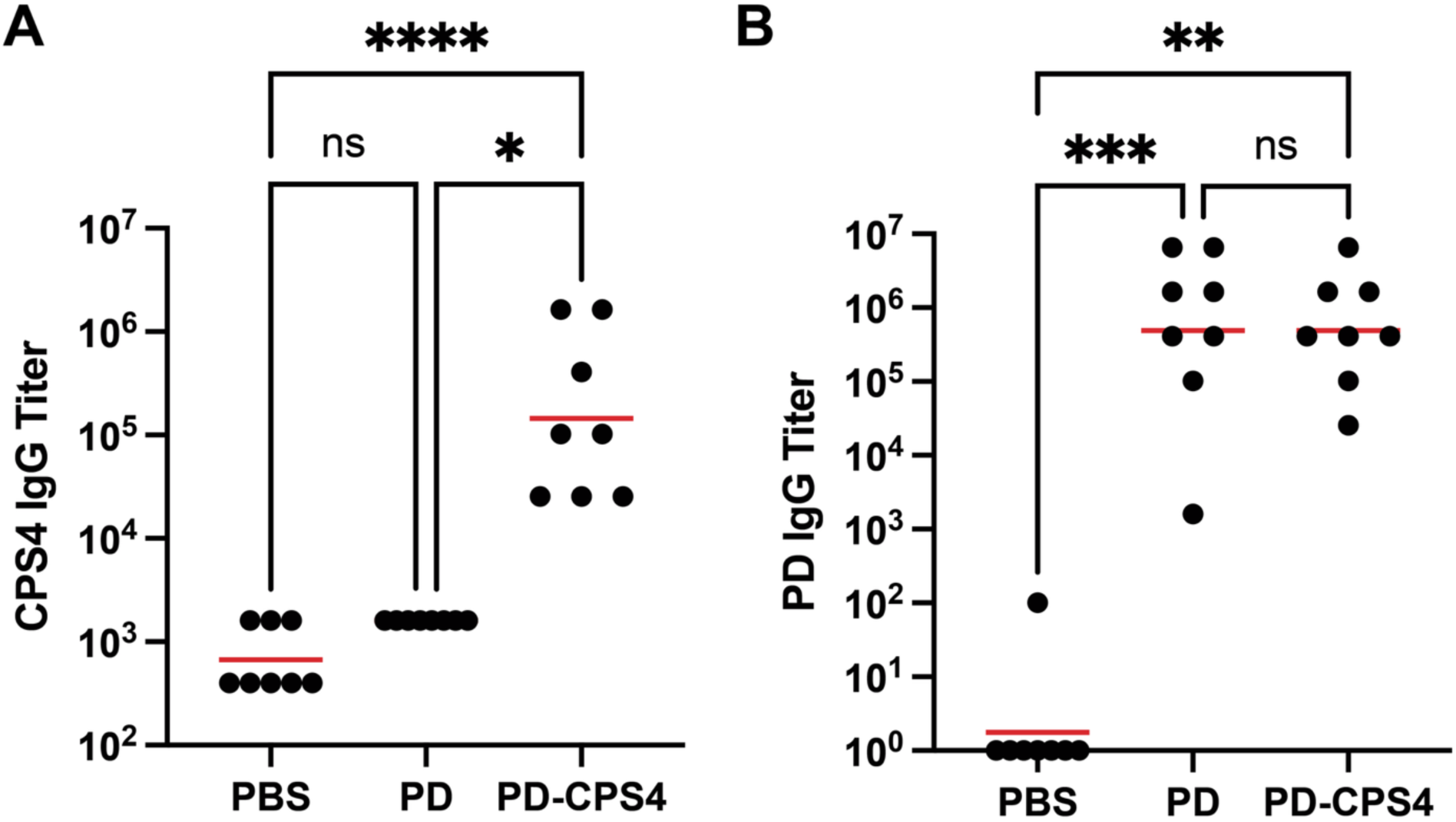
IVG-derived conjugates elicit CPS4-specific antibodies. **(A**) BALB/c mice (*n =* 8 mice per group) were immunized subcutaneously (s.c.) with 10 µg unmodified or CPS4-conjugated PD or PBS as a control. CPS4-specific IgG titers in endpoint (day 77) serum derived from individual mice were determined by ELISA with CPS4 as immobilized antigen. (B) PD-specific IgG titers in same sera from (A) as determined by ELISA with PD as immobilized antigen. Black dots indicate serum IgG titers of individual mice and red lines correspond to mean titers of each group. Statistical significance was determined by the Kruskal-Wallis test with Dunn’s post hoc test for multiple comparisons (\**p* < 0.0332, \*\**p* < 0.0021, \*\*\**p* < 0.0002, and \*\*\*\**p* < 0.0001; ns, not significant).

Carrier protein-specific serum IgG titers were statistically similar between the PD-CPS4 conjugate and unmodified PD carrier, which were both significantly higher than titers measured for mice receiving PBS (**Figure 5B**), confirming the immunogenicity of the carrier protein. Importantly, these findings demonstrate that our *in vitro* platform effectively produced a vaccine candidate capable of generating a robust humoral immune response against a pathogen-specific polysaccharide antigen. This provides an important validation of cell-free synthesized glycoconjugates as rapid and customizable vaccine candidates against bacterial pathogens.

## Discussion

We have optimized the *in vitro* transfer of bacterial glycans onto carrier proteins for improved conjugate vaccine production in cell-free systems. By integrating CFE and glycosylation reactions with a Streptactin-based purification workflow, we were able to generate glycoconjugate vaccines against both ETEC O78 and *S. pneumoniae* CPS4 with greater than 70% glycosylation efficiency and 87% purity (**Table 1**). We also demonstrated that cell-free synthesized and glycosylated conjugate vaccines targeting *S. pneumoniae* CPS4 elicit an anti-glycan immune response in a mouse model.

Compared to our previously developed CFGpS system, coupling CFE with IVG reactions in a two-step process has multiple advantages. First, IVG reactions are modular and can facilitate rapid prototyping efforts for producing diverse vaccine candidates by simply swapping out the glycosylation machinery enriched extract while using the same base CFE reaction for carrier protein expression. Second, the flexible reaction composition of IVG reactions enables the user to optimize for either maximal glycosylation efficiency or total glycoprotein production. Third, by taking advantage of high-yielding extract, IVG reactions could decrease costs per dose by decreasing the amount of costly CFE reagents required to produce the same amount of carrier protein.

Our complete workflow, from CFE expression of carrier protein to *in vitro* glycosylation and subsequent purification, can be performed within five days. Coupling this method with recent advances to increase thermostability and decrease reagent costs for CFE reactions (10) will enable increased preparedness and accessibility to vaccines in low-resource settings or in the event of a pandemic.

## Methods

### DNA preparation

DNA was purchased from Twist Bioscience. Plasmid DNA was prepared using either the ZymoPURE Midi Kit (Zymo Research) or the QIAGEN Plasmid Midi Kit (QIAGEN) according to the manufacturer’s protocols. A list of plasmids is provided in **Table 2**.

**Table 2.** Plasmids used in this study.

| Plasmid | Source |
| --- | --- |
| pSF-CjPglB<sup>Q287K</sup>-LpxE-KanR | (60) |
| pSF-CjPglB-LpxE-KanR | (10) |
| pB-4 | (62) |
| pMW07-O78 | (13, 52, 63) |
| pJL1-PD-4xDQNAT | (13) |
| pJL1-PD-LL-4xDQNAT | This study |

### BL21 (DE3) Star cell extract preparation

Cell-free extracts were prepared as previously described (35, 64, 65). BL21 (DE3) Star strains were inoculated to OD_600_ ≈ 0.06-0.08. When cells reached an OD_600_ ≈ 0.6, 0.5 mM of isopropyl β-d-1-thiogalactopyranoside was added to induce expression of T7 RNA polymerase for use in protein synthesis. Cells were harvested at OD_600_ 3.0 by centrifugation at 8,000 x *g* for 5 min and processed on ice unless otherwise stated. Following centrifugation, the cell pellets were then washed three times in S30 buffer (10 mM Tris acetate, pH 8.2, 14 mM magnesium acetate and 60 mM potassium acetate). Following each wash step, the cells were pelleted at 10,000 x *g* for 2 min and resuspended by vortexing in 15 s increments. Following the last wash steps, the pellets were flash frozen and stored at −80 °C. For lysis, the cells were thawed on ice for 1 h and then resuspended in 1 mL/g S30 buffer and homogenized using an Avestin EmulsiFlex B15 at 20,000 psi using a single pass. The resulting lysate was clarified at 12,000 x *g* for 10 min. The lysate was then centrifuged at 10,000 x *g* for 10 min and the supernatant was flash frozen and stored at −80 °C until further use.

### CLM24 Δ*lpxM* extracts coenriched with *Cj*PglB and ETEC O78 O-PS

For extracts enriched with ETEC O78 O-PS and *Cj*PglB, CLM24 Δ*lpxM* cells were transformed with both pSF-*Cj*PglB-LpxE-KanR and pMW07-O78 plasmids and plated on LB agar containing kanamycin at 50 µg/mL and chloramphenicol at 34 µg/mL. A single transformed colony was then used to grow an overnight culture in LB media supplemented with kanamycin at 50 µg/mL and chloramphenicol at 34 µg/mL. The following day, 2xYTPG media supplemented with kanamycin at 50 µg/mL and chloramphenicol at 34 µg/mL was inoculated at a starting OD_600_ = 0.06-0.08 and incubated at 37 °C with agitation at 250 RPM. At OD_600_ ≈ 0.8, the cells were induced with 0.1% arabinose and the temperature was turned down to 30 °C and agitation set to 220 RPM. Harvest and lysis were performed as described for BL21 (DE3) Star, except that between the two final clarification steps the supernatant was incubated for 1 h at 37 °C with agitation at 250 RPM.

### Hobby extracts coenriched with *Cj*PglB^Q287K^ and CPS from *S. pneumoniae* CPS4

For extracts enriched with *Cj*PglB^Q287K^ and CPS from *S. pneumoniae* CPS4, the above directions provided for extracts enriched with *Cj*PglB^Q287K^ and O-PS from ETEC O78 were followed with the exception that pB-4 rather than pMW07-O78, and pSF-*Cj*PglB^Q287K^-LpxE-KanR rather than pSF-*Cj*PglB-LpxE-KanR, were used for transformation. Additionally, the Hobby strain (61) for increased expression of CPS4 was used rather than CLM24 Δ*lpxM.* For plasmid maintenance, 20 µg/mL tetracyline rather than 50 µg/mL chloramphenicol was used to maintain the glycan synthesis plasmid. To induce for CPS4 overexpression, at OD_600_ = 0.8, in addition to 0.1% arabinose, 0.5 mM isopropyl β-d-1-thiogalactopyranoside was added.

### CLM24 Δ*lpxM* enriched with *Cj*PglB or Hobby extracts enriched with *Cj*PglB^Q287K^

For CLM24 Δ*lpxM* extracts enriched with *Cj*PglB, the above directions provided for extracts enriched with *Cj*PglB and O-PS from ETEC O78 were followed with the exceptions of only pSF-*Cj*PglB-LpxE-KanR was used to transform cells and only 50 µg/mL kanamycin was used to select for and maintain successfully transformed cells. For Hobby extracts enriched with *Cj*PglB^Q287K^, the above directions provided for extracts enriched with *Cj*PglB^Q287K^ and CPS from *S. pneumoniae* CPS 4 were followed with the exceptions of only pSF-*Cj*PglB^Q287K^-LpxE-KanR was used to transform cells and only 50 µg/mL kanamycin was used to select for and maintain successfully transformed cells.

### CFE reactions

CFE reactions were set up as previously described (35, 66, 67). Briefly, reactions were composed of 8 mM magnesium glutamate, 10 mM ammonium glutamate, 130 mM potassium glutamate, 1.2 mM ATP, 0.85 mM of CTP, GTP, and UTP, respectively, 0.034 mg/mL folinic acid, 0.17 mg/mL of E. coli MRE600 tRNA, 0.40 mM NAD, 0.27 mM CoA, 4 mM oxalic acid, 1.5 mM spermidine, 1 mM putrescine, 57 mM HEPES at a pH of 7.2, 30 mM PEP, 2 mM of each amino acid, 13.33 ng/µL plasmid template and 30% v/v *E. coli* extract described above.

Small scale reactions were set up in 200 µL PCR tubes or 2 mL microcentrifuge tubes and incubated for 14-16 h at 30 °C. For larger scale reactions, 250 µL reactions were incubated in 24-well plates (Corning). The plate was covered and incubated at 30 °C at 300 RPM for 14-16 h.

### Cell-free glycoprotein synthesis (CFGpS) reactions

For cell-free glycoprotein synthesis reactions (50), the above directions were followed for CFE reactions with the addition of 0.1 mg/mL T7 RNA polymerase in 50% w/v glycerol and using a concentration of 10 mM magnesium glutamate instead of 8 mM magnesium glutamate. After 2 to 4 h, the reactions were supplemented with a final concentration of 0.1% (w/v) DDM and 25 mM MnCl_2_ to initiate glycosylation and were incubated at 30 °C for an additional 16 h. Before analysis, samples were centrifuged at 16,000 x *g* at 4 °C for 10 min and the soluble fraction was removed.

### *In vitro* glycosylation reactions

To assemble *in vitro* glycosylation reactions, BL21 (DE3) Star CFE reactions expressing a desired carrier protein were mixed with enriched glycosylation extracts and supplemented with a final concentration of 50 mM HEPES pH 7.4, 10 mM McCl_2,_ 1% (w/v) Ficoll 400, 0.1 % (w/v) DDM. Unless otherwise stated, reactions were composed of 10% v/v carrier protein expressing CFE reactions, 80% v/v enriched glycosylation extract, and 10% v/v master mix containing the remaining components required for glycosylation. Before analysis, samples were centrifuged at 10,000 x *g* for 10 min at 4 °C and the soluble fraction was removed.

### Quantification of protein yields in CFE and CFGpS using C^14^ leucine incorporation

Radioactive incorporation was carried out as previously described (67, 68). CFE and CFGpS reactions were assembled according to the above directions with the addition of 10 μM C^14^ leucine (Perkin Elmer). CFGpS reactions were spiked with 0.1% (w/v) DDM and 25 mM MnCl_2_ after 2 to 4 h and incubated overnight at 30 °C while CFE reactions were incubated at 30 °C overnight. Following overnight incubation, reactions were centrifuged at 12,000 x *g* for 10 min at 4 °C. 6 μL of the resulting supernatant was then mixed with 6 μL of 0.5 M KOH and incubated for 20 min at 37 °C. Next, 4 μL of the supernatant-KOH mixture was spotted onto two filtermats (Perkin Elmer) and dried under a heat lamp. Once completely dry, one filtermat was washed three times with 5% w/v TCA for 15 min at 4 °C followed by one wash with ethanol at room temperature for 10 min. After the ethanol wash, the filtermat was then dried under a heat lamp. Scintillation wax (Perkin Elmer) was then melted on both filtermats using a hot plate and radioactive counts were then measured using a MicroBeta2 (Perkin Elmer). Counts from a reaction assembled without template DNA was used as a background subtraction from all reactions. The concentration of leucine incorporated by CFE reactions was then calculated by multiplying the fraction of incorporated radioactive leucine to total radioactive leucine (washed filtermat counts/unwashed filtermat counts) by the total concentration of leucine in the reaction. The amount of protein produced was then determined by dividing the concentration of incorporated leucine by the number of leucine residues in the expressed protein followed by multiplication by the molecular weight of the expressed protein.

### Protein purification

For large scale purifications, proteins were purified as previously described (15). After overnight incubation, IVG samples were centrifuged at 16,000 x *g* for 30 min at 4 °C and the supernatant was transferred into a fresh 15 mL conical tube. Strep-tagged proteins were purified using a 1 mL Strep-tactin XT Sepharose column (Cytiva) with an ÄKTA Avant (Cytiva) according to manufacturer’s protocols. Briefly, the column was equilibrated with 10 CV of Buffer A (PBS pH 7.4, Corning) before the clarified reaction was loaded on to the column followed by 10 mL of Buffer A. To wash the column, 10 CV of Buffer A was added, and elution was performed using a gradient from 0 to 100% Buffer B (PBS, 50 mM biotin; IBA Life Sciences). Elution fractions were collected in 2 mL aliquots at 4 °C in 96-deep well plates (Cytiva).

### Protein dialysis and concentration

Following elution, protein samples were dialyzed into endotoxin free 1x PBS using a Slide-A-Lyzer G3 Dialysis cassette (10K MWCO). Samples were then concentrated using Amicon® Ultra Centrifugal Filter 10 KDa MWCO (Sigma Aldrich) according to the manufacturer’s instructions and sterile filtered using an 0.22 µm syringe filter.

### Endotoxin removal

Endotoxins were removed using Pierce™ High Capacity Endotoxin Removal Resin (Thermo Fisher Scientific) as previously described (15). To regenerate the resin, 2 mL of the resin packed into a 15 mL conical tube was incubated with 10 mL 0.2 N sodium hydroxide for 10 min. The supernatant was removed, and the resin was then incubated with 10 mL 10 N sodium hydroxide at room temperature for 14-16 h. The next morning, the supernatant was removed, and 10 mL 2 M sodium chloride was added and incubated with the resin for 10 min. This 2 M sodium chloride incubation step was then repeated an additional time. Finally, 1 mL of endotoxin free water (Sigma Aldrich) was added to the resin. The regenerated resin was added to a 2 mL Poly-Prep® Chromatography column (Bio-Rad Laboratories) and equilibrated four times with 1 mL of PBS. The pooled purified elution fractions were diluted two-fold in PBS and added to the resin in 1 mL aliquots.

### Endotoxin assay

Endotoxin concentrations were measured using the Pierce™ Chromogenic Endotoxin Quant Kit (Thermo Fisher Scientific) according to the manufacturer’s instructions.

### Purified protein quantification assay

Purified proteins were quantified using the Pierce™ BCA Protein Assay Kit (Thermo Fisher Scientific) according to the manufacturer’s instructions.

### SDS-PAGE protein analysis

Samples were prepared by heating to 100 °C for 10 min in the Invitrogen™ NuPAGE™ LDS Sample Buffer (Thermo Fisher Scientific) supplemented with DTT to a final concentration of 100 mM. Following heat denaturing, the samples were then loaded on a 4-12% Bis-Tris gel and run with MOPS SDS running buffer at 160V until the dye reached the end of the gel. The gels were then stained with Aquastain Protein Gel Stain (BullDog Bio Inc) for 1 h shaking at room temperature, and then destained with Milli-Q water for 1 h shaking at room temperature. Gels were then imaged using a LI-COR Odyssey Fc (LI-COR Biosciences) using the 700-fluorescent channel. Images were analyzed using densitometry on LI-COR Image Studio Lite.

### Western blot analysis

Samples were prepared by heating to 100 °C for 10 min in the Invitrogen™ NuPAGE™ LDS Sample Buffer (Thermo Fisher Scientific) supplemented with DTT to a final concentration of 100 mM. Following heat denaturing, samples were loaded on a 4-12% Bis-Tris gel and run with MOPS SDS running buffer at 100 V for 3 h. Transfer to an 0.45-μm Immobilon-P polyvinylidene difluoride (PVDF) membrane (Sigma Aldrich) was performed using a Trans-Blot® SD Semi-Dry transfer cell (BioRad) according to the manufacturer’s instructions. Membranes were washed in phosphate buffered saline (PBS; Sigma Aldrich) and incubated for 1 h in Intercept (PBS) Blocking Buffer (LI-COR) at room temperature. After blocking, the membrane was washed in PBS for 5 min, three times. Membranes were then probed with either anti-6x-His antibody (Abcam # ab1187, diluted 1:7500), anti-ETEC O78 antibody (Abcam # ab78827. diluted 1:1000), or CPS4 antiserum (Cedarlane, 16747(SS), diluted 1:1000) diluted in Intercept PBS Blocking Buffer supplemented with 0.2% Tween 20 (Sigma-Aldrich), and incubated for 1 h at room temperature. The membrane was washed a total of three times, for 5 min each, in PBS with 0.1% Tween 20. A fluorescent goat, anti-rabbit antibody (LI-COR #GAR680) was used as the secondary antibody at a 1:10,000 dilution in Intercept PBS Blocking Buffer supplemented with 0.2% Tween 20 and 0.1% SDS and incubated for 1 h at room temperature. The membrane was washed a total of three times, for 5 min each, in PBS with 0.1% Tween 20 followed by a single 5-min wash with PBS. Membranes were then imaged using a LI-COR Odyssey Fc and analyzed by densitometry using LI-COR Image Studio Lite.

### Mouse immunization

Groups of eight six-week-old BALB/c mice (Jackson Laboratory) were injected subcutaneously (s.c.) with 50 μL aluminum hydroxide (Thermo Fisher Scientific or InvivoGen) plus 50 μL PBS (pH 7.4) alone or with purified aglycosylated or glycosylated carrier proteins. 10 μg antigen on a total protein basis was used for immunization of all groups. After initial immunizations, boosters of identical doses were given 21, 42, and 63 days later. Blood was collected on days 0, 14, 35, 56, and 77. Mice were observed at 24 and 48 h after each injection for change in behavior and physical health. No abnormal responses were observed. This work was carried out under Protocol 2012-0132 approved by the Cornell University Institutional Animal Care and Use Committee (IACUC).

### Serum antibody titering

To determine *S. pneumoniae* CPS4- and PD-specific IgG titers, sera from immunized mice were subjected to ELISA. Whole blood was centrifuged at 2000 x *g* for 15 min and sera was stored at −20 °C. 96-well MaxiSorp plates (Nunc Nalgene) were coated with 5 μg/mL *S. pneumoniae* CPS Type 4 (ATCC 34-X) in PBS or 2 μg/mL purified PD and incubated overnight at 4 °C. Plates were washed three times with PBST (PBS, 0.05% (v/v) Tween-20, 0.3% (w/v) BSA) and blocked overnight at 4 °C with 5% (w/v) nonfat dry milk in PBS. Sera samples were serially diluted fourfold in duplicate in blocking buffer, from 1:100 to 1:1,638,400 for the *S. pneumoniae* CPS Type 4 ELISA and from 1:100 to 1:26,214,400 for the PD ELISA. Plates were incubated with sera for 2 h at 37 °C. Plates were then washed three times and incubated for 2 h at room temperature with goat anti-Mouse IgG Fc (HRP) (Abcam # ab97265) at 1:25,000 in PBS. After three additional washes with PBST, 1-Step Ultra TMB (3,3′,5,5′-tetramethylbenzidine) ELISA substrate solution (Thermo Fisher Scientific) was added to the plate, which was then incubated at room temperature for 30 min. The reaction was stopped using 2 M H₂SO₄, and absorbance was measured at 450 nm using a microplate spectrophotometer (Molecular Devices). Serum antibody titers were determined by measuring the lowest dilution that resulted in signal 3 standard deviations (SDs) above no serum background controls.

### Statistical analysis

Statistical significance between groups was determined by one-way ANOVA with Tukey’s posthoc test or Kruskal-Wallis (i.e., one-way ANOVA on ranks) with Dunn’s multiple comparisons test (\**p* < 0.0332, \*\**p* < 0.0021, \*\*\**p* < 0.0002, and \*\*\*\**p* < 0.0001; ns, not significant) using GraphPad Prism 9 for MacOS software (version 10).

## Data availability statement

The dataset(s) supporting the conclusions of this article is(are) included within the article and its additional file(s).

## Author contributions

D.A.W., R.A. and S.W.H. designed research, performed research, analyzed data, and wrote the paper. Y.Q., Z.M.S., K.A.M. designed research, performed research, and analyzed data. M.C.J., M.P.D., and A.S.K. designed and directed research, analyzed data, and wrote the paper. All authors read and approved the final manuscript.

## Acknowledgements

This work is supported by the Defense Advanced Research Project Agency (DARPA) contract W911NF-23-2-0039. Any opinions, findings, and conclusion or recommendations expressed in this study are those of the authors and do not necessarily reflect the view of the DARPA or the US government. D.A.W. acknowledges support from the National Science Foundation Graduate Research Fellowship under grant no. DGE-1842165. Z.M.S. acknowledges support from the National Science Foundation National Research Traineeship under grant number 2021900. K.A.M. acknowledges support from the Department of Defense National Science and Engineering Graduate Fellowship number NDSEG11708. We thank Brendan Wren for sharing the Hobby strain.

## Competing interests

M.P.D. and M.C.J. have financial interests in Gauntlet, Inc. and Resilience, Inc. M.P.D. also has financial interests in June Bio APS, Glycobia Inc., UbiquiTx Inc., and Versatope Therapeutics Inc. M.P.D.’s and M.C.J.’s interests are reviewed and managed by Cornell University and Stanford University, respectively, in accordance with their conflict-of-interest policies. All other authors declare no competing interests.

